# Modeling motor neuron resilience in ALS using stem cells

**DOI:** 10.1101/399659

**Authors:** Ilary Allodi, Jik Nijssen, Julio Aguila Benitez, Christoph Schweingruber, Andrea Fuchs, Gillian Bonvicini, Ming Cao, Ole Kiehn, Eva Hedlund

## Abstract

Oculomotor neurons, which regulate eye movement, are resilient to degeneration in the lethal motor neuron disease amyotrophic lateral sclerosis (ALS). It would be highly advantageous if motor neuron resilience could be modeled *in vitro*. Towards this goal, we generated a high proportion of oculomotor neurons from mouse embryonic stem cells through temporal overexpression of Phox2a in neuronal progenitors. We demonstrate, using electrophysiology, immunocytochemistry and RNA sequencing, that *in vitro* generated neurons are *bona fide* oculomotor neurons based on their cellular properties and similarity to their *in vivo* counterpart in rodent and man. We also show that *in vitro* generated oculomotor neurons display a robust activation of survival-promoting Akt signaling and are more resilient to the ALS-like toxicity of kainic acid than spinal motor neurons. Thus, we can generate *bona fide* oculomotor neurons *in vitro* which display a resilience similar to that seen *in vivo*.

## INTRODUCTION

Amyotrophic lateral sclerosis (ALS) is characterized by a loss of motor neurons in cortex, brain stem and spinal cord with subsequent spasticity, muscle atrophy and paralysis. Motor neurons innervating the extraocular muscles, including the oculomotor, trochlear and abducens nuclei, are however relatively resistant to degeneration in ALS (Comley et al., 2016; Gizzi et al., 1992; Nijssen et al., 2017; Nimchinsky et al., 2000). This resilience has been attributed to cell intrinsic properties (Allodi et al., 2016; Brockington et al., 2013; Comley et al., 2015; Hedlund et al., 2010; Kaplan et al., 2014). It would be highly advantageous if differential motor neuron vulnerability could be modeled *in vitro*. This could aid in the identification of gene targets responsible for oculomotor neuron resilience. However, it is not self-evident that *in vitro* generated oculomotor neurons would be more resistant to degeneration in culture than spinal motor neurons, as multiple cell types contribute to motor neuron degeneration in ALS, including e.g. astrocytes (Yamanaka et al., 2008), microglia (Boillee et al., 2006), oligodendrocytes (Kang et al., 2013) and may play a role in selective vulnerability.

During development, distinct neuronal populations are defined through diffusible morphogens with restricted temporal and spatial patterns and subsequent activation of distinct intrinsic transcription factor programs. These programs can be mimicked to specify different CNS regions and neuronal populations from stem cells *in vitro*. When mouse embryonic stem cells (mESCs) are exposed to Shh (sonic hedgehog) signaling and retinoic acid, which ventralize and caudalize the CNS during development, spinal motor neurons constitute 15-30% of the cells in culture, and interneurons the large remainder (Allodi and Hedlund, 2014; Wichterle et al., 2002). Oculomotor neurons constitute only 1% of the cells in the midbrain and thus patterning mESC cultures with Fgf8 and Shh to direct cultures towards a midbrain/hindbrain fate (Allodi and Hedlund, 2014; Hedlund et al., 2008) results in cultures with too few oculomotor neurons to allow for any quantitative analysis. However, when combining morphogens with overexpression of the intrinsic determinant Phox2a in neural precursors a large proportion of Islet1/2+ brain stem motor neurons can be generated (Mong et al., 2014). Furthermore, a combination of the transcription factors Ngn2, Isl1 and Phox2a can specify mESCs into cranial motor neurons (Mazzoni et al., 2013). However, the exact identity of motor neurons generated this way has not been previously demonstrated and it is currently unclear if these are indeed oculomotor neurons or rather a mix of brain stem motor neurons, including for example trigeminal and facial motor neurons.

Towards the goal of developing stem cell-based *in vitro* models of differential vulnerability we generated cultures with a high proportion of oculomotor neurons through temporal overexpression of Phox2a in mESC-derived neuronal progenitors and compared these to mESC-derived spinal motor neurons. Careful characterization of these distinct motor neuron populations using immunocytochemistry, electrophysiological recordings and RNA sequencing demonstrated that mESC-derived oculomotor neurons were highly similar to their *in vivo* counterpart in both rodent and man. Finally, we exposed these to ALS-like conditions using kainic acid and demonstrated that oculomotor neurons were more resilient than spinal motor neurons. Thus, we can generate *bona fide* oculomotor neurons *in vitro* which display a resilience to ALS similar to that seen *in vivo*.

## RESULTS

### *Bona fide* oculomotor neurons can be generated from mESCs

To characterize the specific identity of brain stem motor neurons generated in culture and compare their properties to spinal motor neurons, we generated neurons from mESCs. Spinal motor neurons were derived by adding 100 nM retinoic acid and the smoothened agonist SAG to the culture for five days. Brain stem motor neurons were specified by overexpressing Phox2a under the nestin enhancer and patterning the culture with Fgf8 and SAG for five days (Figure 1A). Immunocytochemical analysis demonstrated that spinal motor neurons expressed the transcription factors Islet1/2 and Hb9 (Mnx1) (64.1±5.2% Islet1/2+Hb9+ cells, mean±SEM, n=4) (Figures 1B, 1D, 1E). Phox2a-overexpressing motor neurons appeared to be of an oculomotor identity as these cells expressed Islet1/2 but lacked Hb9 (6.3±1.3% Islet1/2+Hb9+ cells, mean±SEM, n=4), which is present in all cranial motor neuron nuclei except the oculomotor nucleus (Figures 1C, 1D, 1F, 1G) (Guidato et al., 2003; Lance-Jones et al., 2012). From hereon we therefore refer to the brain stem motor neurons as oculomotor neurons. In oculomotor neuron cultures 47.5±5.9% (mean±SEM, n=4) of βIII-tubulin+ cells were Islet1/2+ and 62±5.2% (mean±SEM, n=4) of the Islet1/2+ cells were Phox2a+, which demonstrates that 25% of all neurons generated were oculomotor neurons (Figure 1G).

**Figure 1.**
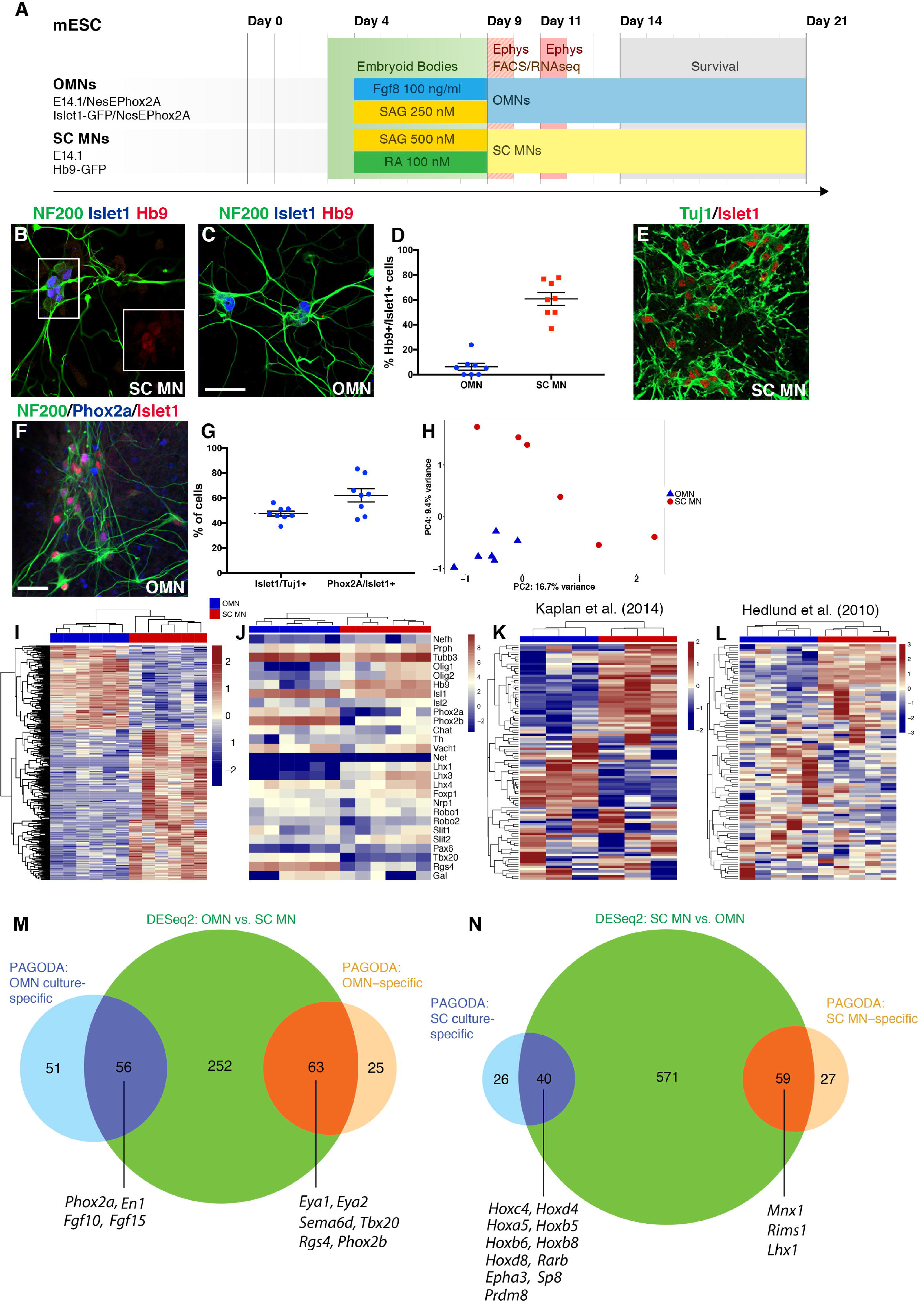
Phox2a overexpression drives the generation of *bona fide* oculomotor neurons from stem cells. (A) Time line depicting in vitro protocols for resistant oculomotor neurons (OMN) and vulnerable spinal motor neuron (SC MN) generation. ESC column reports the four cell lines used in this study. Patterning was performed for 5 days after ESC expansion and EB formation. mRNA-seq coupled with FACS was performed on EB dissociation day (day 9 of the protocol). Survival assay was performed after five days from EB dissociation. (B-C) SC MNs generated from E14.1 ESCs express Hb9, Islet1 and NF200, OMNs generated from E14.1 ESCs over-expressing the transcription factor Phox2a under the Nestin enhancer co-express Islet1 and NF200 in absence of Hb9. Scale bar in c 60 μm. (D) Percentage of Hb9+/Islet1+ cells in SC MN (64.1±5.2%, mean±SEM, n=4) and OMN (6.3±1.3%, mean±SEM) cultures respectively. (For quantification, experiments were run in quadruplicates including two technical replicates per experiment, with at least 120 Islet1+ cells counted per condition). (E) Microphotographs showing Islet1+/Tuj1+ cells in OMN cultures, (F) OMNs also express the specific marker Phox2a as indicated by asterisks. Scale bars in e and g 100 μm. (G) Quantification of Islet1+ over Tuj1+ cells demonstrates that half neuronal population appears to be Islet1+ (47.5±5.9%, mean±SEM, experiments performed in quadruplicates with two technical replicates each, total number of Tuj1+ cells counted: 1325). Quantification of Phox2A+ over Islet1+ cells (experiments performed in quadruplicates with two technical replicates each, total number Islet1+ cells counted: 746) indicates that 62±5.2% (mean±SEM) of the Islet1+ population is also Phox2a+. All quantifications were performed 5 days after EB dissociation (mean ± SEM). (H) PCA of OMN and SC MN samples based on all genes expressed confirmed cell differential identities. mRNA-seq analysis of OMNs and SC MNs isolated by FACS was performed after 5 days of patterning (day 9 of the protocol). (I) 1,017 differentially expressed genes were found between the two different cell types (adjusted P<0.05). Heatmap shows the top 500 most significant differentially expressed genes by adjusted p-value. (J) Heatmap of expected progenitors and motor neuron transcripts but also OMN and SC MN specific transcripts obtained in the two generated motor neuron populations. (K) Using the top 100 DEGs obtained from our RNAseq analysis we could separate datasets originating from *in vivo* microarray studies on early postnatal (J) and adult (L) rodent OMNs and SC MNs. (M) Venn diagram showing gene sets enriched either in brain stem cultures or OMN-specific as revealed by PAGODA analysis, (N) gene sets preferentially found in spinal cord cultures or MN-specific.

Oculomotor neurons (derived from Islet-GFP mESCs) and spinal motor neurons motor neurons (derived from Hb9-GFP mESCs) were analyzed for their electrophysiological properties at day 9 and day 11 time points (Figure S1). Cells were able to fire overshooting action potentials in response to depolarizing current pulses. At day 9 few spikes were evoked. Neuronal maturation from day 9 to 11 was seen as change in the electrical properties where day 11 oculomotor neurons could fire trains of action potentials (Figure S1A and S1B) and the width of action potentials was decreased when compared with day 9 (Figure S1C). The increase in the size of neurons was reflected by the increased cell capacity (Figure S1D), decreased input resistance (Figure S1E) and increased rheobase (Figure S1F) from day 9 to 11. Similar changes were seen in spinal motor neurons, which fired a higher number of action potentials (Figure S1G, H) and showed a reduced width of action potentials (Figure S1I) at day 11 compared to day 9 time point. Moreover, the changes in cell capacity (Figure S1J), input resistance (Figure S1K) and rheobase (Figure S1L) showed signs of neuronal maturation over time. To further investigate neuronal maturation over time, ChAT staining was performed on oculomotor and spinal motor neuron cultures confirming expression of the transferase from day 14 *in vitro* in both Islet1+/Hb9- and Islet+/Hb9+ cells (Figures S1M and S1N).

To further confirm the identity of the cells generated we sorted the motor neurons based on GFP expression using FACS. For this purpose we used a Hb9-GFP mESC line to select for spinal motor neurons and an Islet-GFP::NesEPhox2a mESCs line for the purification of oculomotor neurons (Figure 1A and Figures S2A, S2B, S2C and S2D). PolyA-based RNA sequencing of the sorted fractions revealed a similar number of detected genes for both lines (RPKM > 0.1 OMN 13941±293.5 and SC MN 14660.3±304.1; RPKM > 1 OMN 10507±296 and SC MN 10914.3±362.7 mean ± SEM, t test for RPKM > 0.1 *P*=0.1196, t test for RPKM > 1 *P*=0.4047, Figure S2E).

Sorted oculomotor neurons clustered closely together in the principal component analysis (PCA) and were separated from spinal motor neurons along the fourth principal component (Figure 1H). Hierarchical clustering of the 1,017 genes that were differentially expressed between oculomotor neurons and spinal motor neurons at an adjusted *P* value <0.05 demonstrated that each group is defined by a specific set of markers, with a majority of genes being more highly expressed in spinal motor neurons (Figure 1I and Table S1). Analysis of Hox mRNAs clearly demonstrated the more caudal nature of spinal motor neurons compared to oculomotor neurons (Figure S1F). The oculomotor neurons and spinal motor neurons clustered separately based on known marker gene expression e.g. *Phox2a*, *Phox2b*, *Tbx20* and *Rgs4* for oculomotor neurons and *Olig1/2*, *Hb9* and *Lhx1/3* for spinal motor neurons. Oculomotor neuron cultures were also devoid of noradrenergic neuron markers *Slc6a2* (*NET*) and *Th* (Figure 1J).

Using the top 100 differentially expressed genes (DEGs) between our *in vitro* generated oculomotor neurons and spinal motor neurons, we could separate a published microarray data set (Mazzoni et al., 2013) on *in vitro* generated brain stem and spinal motor neurons (Figure S1G). Importantly, we could also use the 100 top DEGs between our oculomotor and spinal motor neurons to separate microarray data originating from mouse oculomotor and spinal motor neurons isolated from P7 mice (Kaplan et al., 2014) (Figure 1K) and from adult rats (Hedlund et al., 2010) (Figure 1L). Thus, our *in vitro* generated motor neuron subpopulations closely resemble their *in vivo* counterparts.

To further delineate the nature of our *in vitro* culture systems, we set out to identify gene clusters regulated in groups of samples in an unbiased way. Here we analyzed all GFP-positive (Hb9+ for spinal and Islet1+ for oculomotor) and -negative fractions from both spinal- and midbrain-specified cultures using PAGODA (Fan et al., 2016). This analysis revealed gene sets enriched in the spinal and oculomotor cultures (GFP+ and GFP-samples), as well as in the motor neuron groups specifically (GFP+) (Figure 1M and 1N, Table S2). Of the oculomotor-enriched genes found with DESeq2, a subset was enriched in all midbrain-specified cultures (GFP+ and GFP-cells), such as *Phox2a* and Engrailed1 (*En1*). *Fgf10*, known to be expressed in the developing midbrain, including the oculomotor nucleus (Hattori et al., 1997) and Fgf15, with known enrichment in midbrain and rhombomere 1 (Partanen, 2007) were also enriched in our midbrain-specified cultures (Figure 1M). The genes that were highly enriched specifically in oculomotor neurons (GFP+) included *Eya1*, *Eya2*, *Tbx20* and *Phox2b* (Figure 1M). Vice versa, of the genes found to be differentially expressed in spinal motor neurons versus oculomotor neurons with DESeq2, a subset was enriched in all samples of the spinal-specified cultures versus the midbrain-specified cultures. This included Hox genes which regulate positional identity e.g. *Hoxc4*, *Hoxd4*, *Hoxa5*, *Hoxb5*, *Hoxb6*, *Hoxb8* and *Hoxd8*, as well as spinal V1 interneuron markers *Sp8* and *Prdm8*. Another subset was found to be highly specific for only the spinal motor neurons, such as *Mnx1* (Hb9) and *Lhx1* (Figure 1N).

Different axon guidance-related genes are expressed by oculomotor and spinal cord motor neurons during development (Bjorke et al., 2016; Gutekunst and Gross, 2014; Stark et al., 2015; Wang et al., 2011). To further characterize the *in vitro* generated oculomotor neurons, we analyzed their mRNA expression of morphological markers that are important for axon guidance. We found that oculomotor neurons were enriched in *Plxna4, Sema6d*, *Cdh6*, *Cdh12*, while spinal motor neurons were enriched for *Epha3, Ephx4, Sema4a* and *Sema5b* (Figure S2G). We subsequently analyzed the expression level of these axon guidance RNAs in a microarray data from mouse oculomotor and spinal motor neurons isolated from P7 mice (Kaplan et al., 2014) and found that Sema6d was still enriched in oculomotor postnatally (Figure S2H).

Thus, our *in vitro* generated oculomotor neurons have characteristics distinct from spinal motor neurons regarding genes that govern axon guidance and some of these characteristic are maintained *in vivo*.

In conclusion, we have generated a high proportion of *bona fide* oculomotor neurons *in vitro* that can be utilized to understand their normal function as well as resilience to degeneration in motor neuron diseases.

### Stem cell-derived oculomotor neurons are relatively resistant to excitotoxicity

To investigate if *in vitro* generated oculomotor neurons are relatively resistant to ALS-like toxicity similarly to their *in vivo* counterparts, we used kainic acid which is a neuro-excitatory amino acid that acts by activating glutamate receptors. Glutamate excitotoxicity is thought to be a downstream event in motor neuron degeneration in ALS and thus considered appropriate to model this disease. First, we evaluated the mRNA levels of glutamate ionotropic receptors in our cultures to ensure that generated neurons could respond to kainic acid. mRNA sequencing data of sorted motor neurons demonstrated that oculomotor neurons and spinal motor neurons expressed similar levels of AMPA, NMDA and kainate receptor subunits (Figure 2A). Grik5 (glutamate receptor, ionotrophic kainate 5) was the subunit expressed at highest level (Figure 2A) and it was also detectable at the protein level in both oculomotor (Figure 2B) and spinal (Figure 2C) motor neurons. Thus, both motor neuron types should respond to kainic acid. Exposure of the cultures to kainic acid for one week demonstrated that oculomotor neurons were more resilient to the elicited long-term excitotoxicity than spinal motor neurons (ANOVA, \**P* < 0.05, Figure 2D-F).

**Figure 2.**
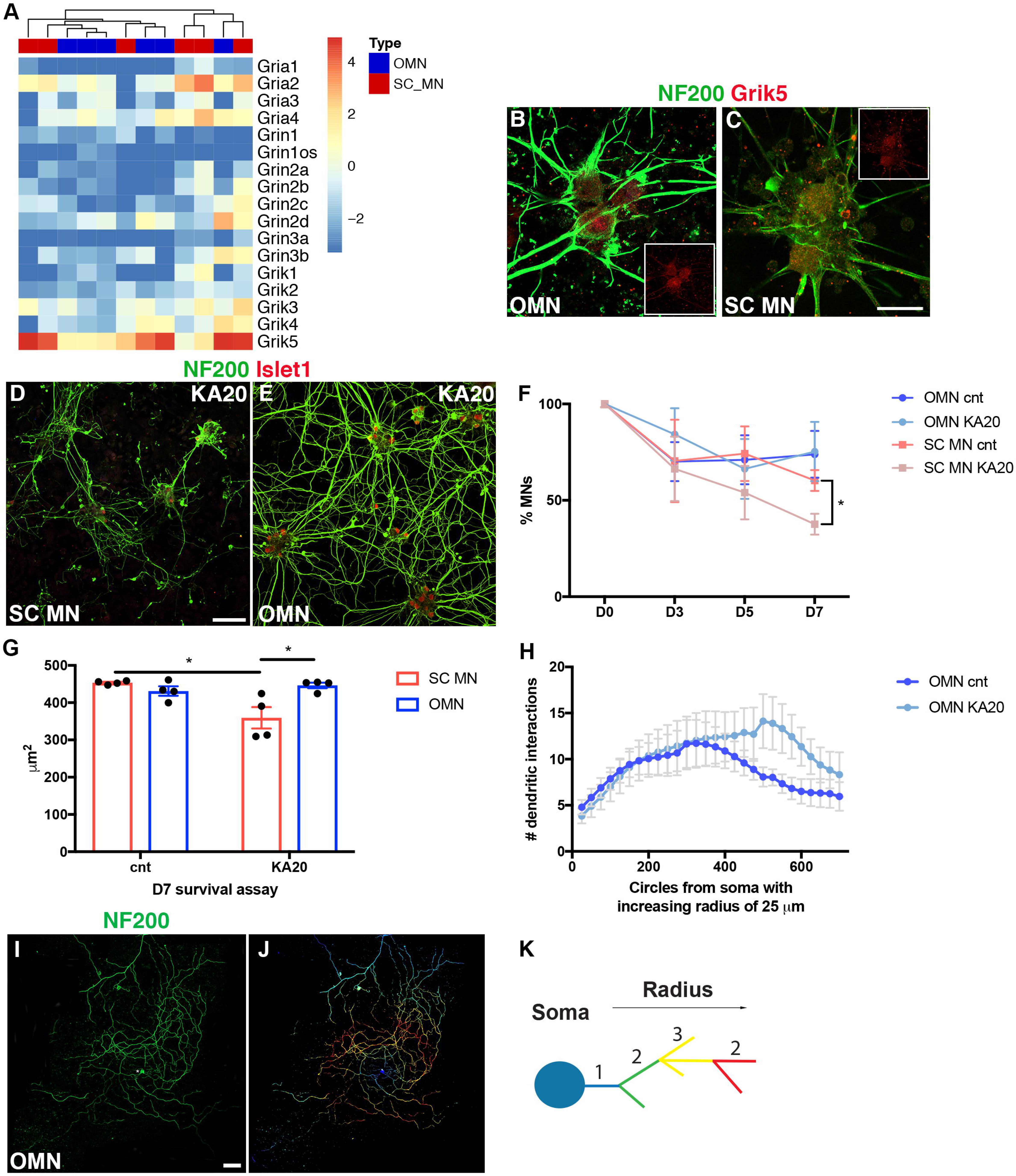
*In vitro* generated oculomotor neurons are relatively resistant to ALS-like toxicity. (A) Oculomotor neurons (OMNs) and spinal motor neurons (SC MNs) express similar mRNA levels of glutamate ionotropic receptor AMPA, NMDA and kainate type subunits. The heatmap shows log2 RPKM values of these subunits and no separate clustering of the two cell types is observed. (B-C) Immunohistochemistry performed on generated OMNs and SC MNs at D1 in vitro in control conditions; similar levels of glutamate ionotropic receptor kainate type subunit 5 (Grik5) are found in both cell types. Scale bar in d 60 μm. (D-E) Microphotographs presenting SC MN and OMN response to kainic acid induced toxicity (20 μM) for a week. Scale bars in f = 100 μm. (F) Curves represent percentages of MN survival over time in OMN and SC MN cultures. OMNs were visualized as NF200+Islet1+Hb9- clls, while SC MNs as NF200+Islet1+Hb9+ cells. OMNs show increased survival to KA toxicity at D7 (mean ± SEM, 2way ANOVA and Tukey’s multiple comparison test, F(9, 56)=2.333, *P=0.0261, SC MNs n=4; OMNs n=5) when compared to SC MNs (experiments were performed at least in quadruplicates, with technical replicates and with at least 130 motor neurons counted per condition in each experiment). Analysis of the length of neuronal processes in both oculomotor and spinal motor neuron cultures exposed to kainic acid for seven days. showed that oculomotor neurons were unaffected by kainic acid while spinal motor neurons displayed a shortening of neurites (G). (H-K) Sholl analysis was performed on OMN at D7 survival assay in control and KA20 conditions to further assess individual MN arborization complexity during toxicity. Scale bar = 100 μm. (I) Sholl mask was applied to individual OMNs after specifying the radius from the center of the soma of the neuron and created concentric circles every 25 μm of increasing radius. (J) Comparison of average number of neurite intersections of OMN in control and KA20 toxicity conditions with radial step size of 25 μm. OMNs did not show reduction in arborization (multiple t test, n = 10 per condition). (K) Schematic depicting identification of neurite segments by Sholl analysis. Color code is assigned depending on arbor localization from the soma in an inside-out manner following the given radius. Multiple intersections within the same segment display the same colour.

Stem cell-derived cultures are not temporally restricted in a precise way. Cultures that contain post mitotic neurons will also harbor neuronal progenitors that later on are specified into neurons. It can thus be challenging to distinguish survival from renewal in a stem cell-based *in vitro* system. We therefore analyzed the turn-over rate in our cultures by pulsing these with BrdU. Analysis of the number of BrdU+ Islet1/2+ neurons demonstrated that a significantly higher number of motor neurons were generated over time in the spinal motor neuron cultures compared to the oculomotor neuron cultures (Figure S2I). Consequently, the resilience of oculomotor neurons should be considered even more pronounced as these cultures had a comparatively low turn-over rate. Furthermore, we investigated the presence of astrocytes in the cultures and found that only a minority of cells expressed GFAP (Figure S2J and S2K). As the processes of motor neurons show the first signs of pathology in ALS we investigated length of neuronal processes in both oculomotor and spinal motor neuron cultures exposed to kainic acid for seven days. The result clearly demonstrated that oculomotor neurons were unaffected by kainic acid while spinal motor neurons displayed a shortening of neurites (Figure 2G). To further evaluate oculomotor neurites, we used Sholl analysis to interrogate arborization complexity. The number and branching of oculomotor neurites were not diminished under kainic acid treatment (Figure 2H-K). As axonal fragmentation and degeneration is seen early on in motor neurons in ALS, our data clearly demonstrates that oculomotor neurons cope well under conditions of excitotoxicity and maintain normal morphology.

In conclusion, *in vitro* generated oculomotor neurons are highly resilient to ALS-like excitotoxicity. Thus, the pattern of selective motor neuron vulnerability seen *in vivo* can be replicated *in vitro*.

### Oculomotor neurons have high levels of calcium buffering proteins and Akt signaling which could contribute to resilience

To explain the resilience of *in vitro* generated oculomotor neurons to excitotoxicity we conducted a directed comparative analysis of the RNA sequencing data from purified oculomotor neurons versus spinal motor neurons. It has been suggested that preferential expression of calcium binding proteins in oculomotor neurons plays a role in their resilience (Comley et al., 2015; Van Den Bosch et al., 2002). We focused on transcripts with implications in calcium handling. Our analysis demonstrated that Cald1, Esyt1, Camk2a and Hpca11 were preferentially expressed in oculomotor neurons (Figure S3A). Esyt1 protein levels were also preferential to oculomotor neurons and unaffected by kainic acid treatment (Figure S3B, S3C and S3D). This could render oculomotor neurons with an increased capacity to buffer Ca^2+^ intracellularly.

We have previously shown that Akt signaling is important in motor neuron resilience towards ALS after IGF-2 treatment (Allodi et al., 2016). We therefore analyzed our RNA sequencing data from sorted oculomotor and spinal motor neurons for Akt signaling effectors which could aid in explaining the relative resilience of oculomotor neurons *in vitro*. This analysis demonstrated an elevated expression of Akt1 and Akt3 in oculomotor neurons compared to spinal motor neurons (adjusted *P* < 0.05, Figure 3A). Consistent with our RNA sequencing findings, immunocytochemistry against pAkt and quantification of fluorescent intensities in Islet1/2^+^ oculomotor and spinal motor neurons derived from mESCs demonstrated that oculomotor neurons had higher levels of pAkt protein than spinal motor neurons (Figure 3B, 3C and 3D). These levels were maintained after excitotoxicity elicited by kainic acid (2-way ANOVA, *** *P* < 0.0001, Figure 3D).

**Figure 3.**
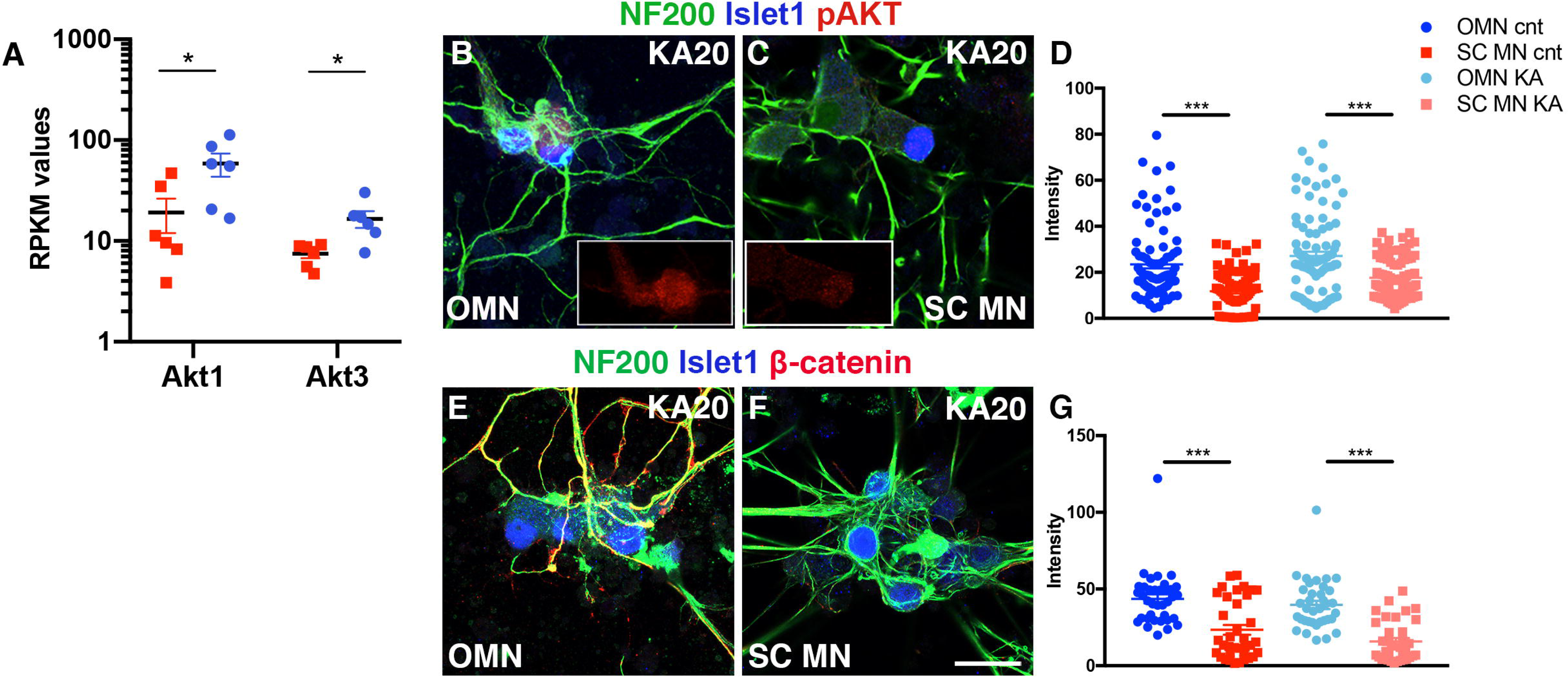
mESC-derived oculomotor neurons have high levels of Akt signaling. (A) RPKM values from RNA sequencing of mESC-derived oculomotor and spinal motor neurons enriched by FACS, shows preferential expression of Akt1 and Akt3 transcripts in the generated oculomotor neurons (OMNs) (adjusted *P < 0.05). (B-C) Microphotographs show pAkt levels in oculomotor (OMN) and spinal motor neuron (SC MN) cultures at D7 toxicity assay assessed by immunohistochemistry, pAKT staining alone in small inserts (red channel). (D) Fluorescense intensities, analyzed in a semi-quantitative manner, indicate higher expression of pAkt in OMNs (mean ± SEM, 2way ANOVA and Tukey’s multiple comparison test, F(1, 353)=66.1, *** P < 0.0001, n=4). (E-G) β-catenin expression is maintained in OMN also during kainate toxicity. Scale bar = 60 μm. (G) Semi-quantitative analysis of β-catenin over NF200 staining intensities in OMNs and SC MNs in kainate conditions (mean ± SEM, t test, t=6.799 df=69, *** P < 0.0001, n=3). Scale bars = 60 μm.

To further validate this pathway in oculomotor neuron resistance we analyzed the protein levels of activated β-catenin (non-phosphorylated), a down-stream effector of Akt signaling, using immunocytochemistry (Figure 3E and 3F). The semi-quantitative analysis demonstrated that oculomotor neurons had elevated levels of β-catenin compared to spinal cord motor neurons (2-way ANOVA, *** *P* < 0.0001, Figure 3G). β-catenin expression was maintained at relatively high levels in oculomotor neurons compared to spinal motor neurons also during kainate toxicity, with expression preferentially localized to processes (Figure 3F and 3G).

Our data thus indicate that Akt signaling, which is a known survival pathway, could contribute to oculomotor neuron resistance in mouse *in vivo* as well as in a dish.

### AKT signaling is elevated in human post mortem oculomotor neurons

To evaluate if our *in vitro* data recapitulates motor neuron resistance in human, we used the spatial transcriptomics method LCM-seq (Nichterwitz et al., 2018; Nichterwitz et al., 2016) to analyze inidividually isolated oculomotor neurons (Figure S4A, S4B, S4C and S4D) and spinal motor neurons (cervical and lumbar spinal cord) (Figure S4E, S4F, S4G and S4H) from human *post mortem* tissues. In addition, we also isolated motor neurons from the Onuf’s nucleus in the sacral spinal cord (Figure S4I, S4J, S4K and S4L and Table S3), as these are highly resilient to degeneration in ALS and maintained until end-stage of disease (Mannen et al., 1977), similar to oculomotor neurons. Analysis of marker gene expression showed that all motor neuron groups expressed high levels of neurofilaments and *VACHT* (*SLC18A3*) (Figure S4M). Motor neurons also expressed *ISL1/2* and *CHAT*, while being almost devoid of contaminating glial markers e.g. *SLC1A3*, *AQP4*, *CCL3*, *AIF1*, *SOX10*, and *MOG* (Figure S4M). Analysis of the *PHOX2A/B* and *HOX* gene code expression clearly clustered oculomotor neurons away from all spinal motor neuron groups (Figure S4N). Principal component analysis (PCA) of all expressed genes separated oculomotor neurons and spinal motor neurons along PC2, while Onuf’s nucleus motor neurons separated out on PC3 (Figure 4A). Analysis of the PI3K-AKT signaling pathway (Figure S4O) demonstrated that *AKT3* was elevated in human oculomotor neurons (adjusted *P* < 0.05, Figure 4B), similar to mESC-derived oculomotor neurons (Figure 3A). To analyze if oculomotor and Onuf’s motor neurons shared gene expression that was distinct from other spinal motor neurons (cervical and lumbar) we analyzed DEGs of oculomotor versus spinal and Onuf’s versus spinal (Table S4 and S6). This analysis showed that the majority of DEGs were unique to each cell type, with 1,025 DEGs unique to oculomotor (553 up- and 472 down-regulated compared to other spinal motor neurons) and 921 DEGs unique to Onuf’s motor neurons (349 up- and 572 down-regulated compared to other spinal motor neurons) (Figure 4C and Table S5 and S6). High *AKT3* levels were unique to oculomotor neurons. Elevated *PVALB* (Parvalbumin) and *GABRA1* levels were also unique to oculomotor neurons (Figure 4C), consistent with previous findings (Brockington et al., 2013; Comley et al., 2015; Hedlund et al., 2010). Oculomotor and Onuf’s shared 214 DEGs, of which 58 were up-regulated and 142 down-regulated compared to other spinal motor neurons, and 14 which were regulated in the opposite direction in the two nuclei (Figure 4C and Table S6). *MIF* (macrophage migration inhibitory factor), which is neuroprotective in ALS (Shvil et al., 2018) showed shared elevated expression in both oculomotor and Onuf’s motor neurons (Figure 4C). GO term analysis of the identified DEGs revealed a number of GO terms that were up-regulated in oculomotor neurons versus the other motor neuron groups, including gluconeogenesis, regulation of membrane potential, anterograde trans-synaptic signaling, cation transmembrane transport, neurodevelopment, neuron differentiation and central nervous system development (Figure 4D and Table S7). Analysis of GO terms enriched in Onuf’s motor neurons included synaptic signalig, anterograde, trans-synaptic signlaing, NAD biosynthetic process, ATP biosynthetic process, regulated exocytosis, regulation of neurotransmitter levels and synaptic vesicle localization (Figure 4E and Table S8).

**Figure 4.**
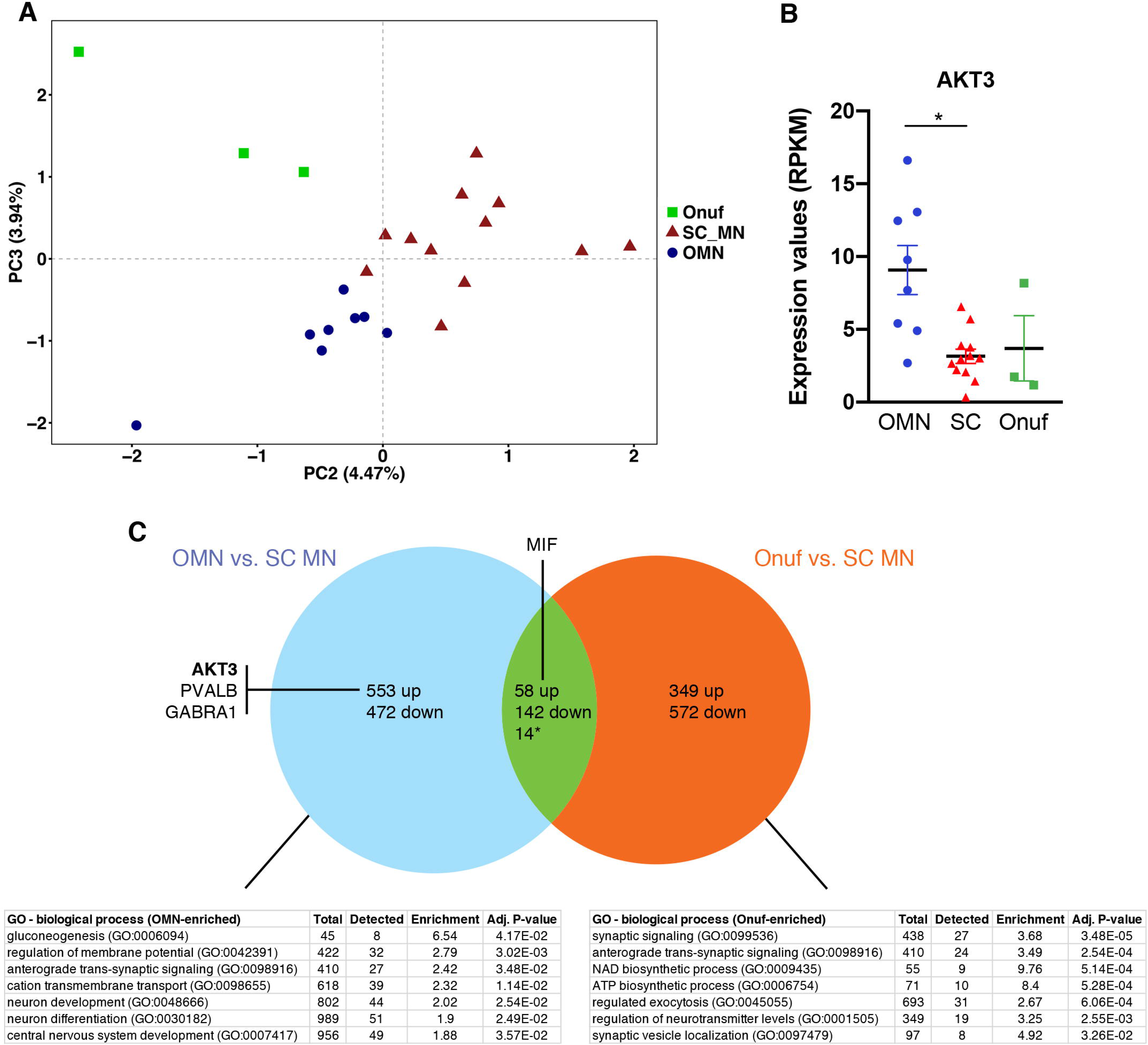
Human oculomotor neurons show a transcriptional profile, with elevated AKT3, unique from Onuf’s nucleus motor neurons. (A) Principal component analysis (PCA) of all expressed genes separated Onuf’s, oculomotor (OMN) and spinal (cervical and lumbar) (SC_MN) motor neuron groups. (B) RNA sequencing data showed that AKT3 was elevated in OMNs compared to the other motor neuron groups (adjusted *P* <0.05). (C) To distinguish the number of DEGs that were shared or unique to each resistant neuron group compared to vunerable spinal motor neurons we analyzed DEGs between OMN and SC_MN and between Onuf’s and SC_MN. This analysis showed that the majority of DEGs were unique to each resistant motor neuron group, with OMNs showing 553 up- and 472 down-regulated unique DEGs and Onuf’s motor neurons showing 349 up- and 572 down-regulated unique DEGs. Oculomotor and Onuf’s shared 58 up-regulated and 142 down-regulated DEGs compared to vulnerable spinal motor neurons. * 14 DEGs were regulated in the opposite direction in the two motor nuclei.

In conclusion, our data indicates that AKT signaling is elevated also in adult human oculomotor neurons and may in part underlie their resilience to degeneration in ALS. Notably, while oculomotor and Onuf’s nucleus motor neurons mainly show distinct gene regulation we also identified common pathways that may underlie their joint resilience.

## DISCUSSION

In ALS, motor neuron subpopulations show differential vulnerability to degeneration. In particular, cranial motor neurons of the oculomotor, trochlear and abducens (cranial nerves 3, 4 and 6), which innervate the extraocular muscles around the eyes are highly resistant throughout disease progression. In our study we demonstrate that a high proportion of *bona fide* oculomotor neurons can be generated from mESC through temporal over-expression of Phox2a, and that these are relatively resilient to ALS-like toxicity.

Oculomotor neurons are generated in the ventral midbrain through the specification by Shh (secreted from the floor plate and the notochord) Fgf8 and Wnts (both secreted by and around the isthmic organiser). These morphogens in turn regulate transcription factors that influence differentiation of ventral midbrain neurons, including oculomotor neurons, dopamine neurons, red nucleus neurons and GABA-positive interneurons (reviewed in (Nijssen et al., 2017)). Oculomotor neurons are specified by the transcription factors Phox2a, Phox2b and Lmx1b (Deng et al., 2011; Pattyn et al., 1997). Phox2a is required to drive oculomotor neuron fate as demonstrated by the lack of both oculomotor and trochlear nuclei in Phox2a knockout mice (Pattyn et al., 1997). Phox2a is also sufficient to generate a complete oculomotor complex as shown by studies in chick (Hasan et al., 2010; Pattyn et al., 1997). Phox2b on the other hand is sufficient to induce ectopic generation of oculomotor neurons in the spinal cord, but it is not required to induce oculomotor neuron specification in the midbrain (Dubreuil et al., 2000; Pattyn et al., 1997). *In vitro* studies have shown that; *i)* mESCs can be directly programmed into cranial neurons using the proneuronal gene *Ngn2*, in combination with *Islet1* and *Phox2a* (Mazzoni et al., 2013), and that *ii)* over-expression of either Phox2a or Phox2b alone in mESC-derived neural progenitors exposed to Shh and Fgf8 can promote a midbrain/hindbrain motor neuron fate (Mong et al., 2014). However, while it was demonstrated that cranial motor neurons were generated *in vitro* it was never previously determined if oculomotor neurons were specifically produced.

Here, we show that Phox2a overexpression in neural progenitors in combination with patterning using Fgf8 and Shh results in the generation of 50% Islet+ neurons of which more than half were Phox2a+. The transcriptome and proteome of oculomotor neurons is distinct from other motor neurons (reviewed in (Nijssen et al., 2017)). Notably, oculomotor neurons can be distinguished by their lack of the transcription factor Hb9, which defines other somatic motor neurons (Guidato et al., 2003; Lance-Jones et al., 2012). As the vast majority of Islet+ cells in our midbrain cultures lacked Hb9 we conclude that the Phox2a+ cells generated were indeed oculomotor neurons and that Phox2a overexpression is sufficient to induce an oculomotor fate from stem cells *in vitro*. This conclusion was further supported by our RNA sequencing experiments on purified oculomotor and spinal motor neuron cultures and our bioinformatics cross-comparison with other *in vitro* and *in vivo* data set from rodent and man (Brockington et al., 2013; Hedlund et al., 2010; Kaplan et al., 2014), which confirmed that the oculomotor neurons we generated were molecularly similar to their *in vivo* counterpart. Furthermore, our analysis of axon guidance molecules demonstrated differential expression of distinct morphological markers in oculomotor and spinal motor neurons. Finally, the electrical properties of the mESC-derived motor neurons matured over time in culture, and cells also expressed the cholingeric marker ChAT. Altogether this demonstrates that we can generate *bona fide* oculomotor neurons.

Furthermore, we found that mRNAs of Fgf10 and Fgf15 were enriched in our oculomotor cultures. These two morphogens are normally expressed in the developing midbrain and Fgf10 is even present within oculomotor neurons themselves (Hattori et al., 1997; Partanen, 2007). It is therefore highly conceivable that these factors are involved in further specifying the cultures. We therefore believe that these factors can be used to further improve the *in vitro* differentiation protocol, but this remains to be further investigated.

Multiple studies have demonstrated that differential motor neuron vulnerability appears to be regulated by cell intrinsic differences in gene expression (Allodi et al., 2016; Brockington et al., 2013; Hedlund et al., 2010). We therefore hypothesized that *in vitro* generated oculomotor neurons would be more resilient to ALS-like toxicity than spinal motor neurons. Indeed, our experiments on neuronal vulnerability clearly demonstrated that oculomotor neurons were more resistant to the ALS-like toxicity elicited by kainic acid than spinal motor neurons. Furthermore, the fact that oculomotor neuron processes remained intact indicates that these cells were indeed highly resilient as axonal degeneration is an early sign of pathology in ALS.

It has been shown that oculomotor neurons have a higher calcium buffering capacity than other motor neuron groups (Vanselow and Keller, 2000), which could render the cells more resilient to excitotoxic insuts. Our finding that several transcripts with implications for calcium handling were preferentially expressed in stem cell derived oculomotor neurons and also in oculomotor neurons in man were thus very compelling and consistent with their resilience. Esyt1, for example, is an ER integral membrane protein which presents a cytosolic synaptotagmin-like domain that can bind organelle membranes and a C2 domain able to bind the plasma membrane. These interactions are involved in exchange of lipids and vescicles but also in the ER control of Ca^2+^ homeostasis. Thus, ER-plasma membrane junctions can control Ca^2+^ release from ER in response to extracellular Ca^2+^ levels (Giordano et al., 2013), an event that could have a beneficial effect during excitotoxicity.

Furthermore, we have previously shown that Akt signaling plays an important role in induced resistance in vulnerable motor neurons (Allodi et al., 2016). We now demonstrate that Akt signaling is more highly activated in resistant oculomotor neurons *in vitro* as well as *in vivo* in human oculomotor neurons. This observation indicates that oculomotor neurons preferentially activate a cell intrinsic survival program. Notably, cross-comparison of human oculomotor neurons with resilient human Onuf’s nucleus motor neurons showed that elevated AKT signaling was unique to oculomotor neurons. When comparing human oculomotor and Onuf’s motor neurons with that of vulnerable spinal motor neurons it became evident that the majority of DEGs were unique to each resilient nucleus. However, there were commonly regulated genes across these two resilient nuclei, including MIF. MIF was recently shown to inhibit the toxicity of misfolded SOD1 (Shvil et al., 2018), indicating that these two neuron groups may be better at handling toxic aggregation-prone proteins. In conclusion, we have demonstrated that *bona fide* oculomotor neurons can be generated from stem cells *in vitro.* We also show that these neurons are relatively resilient to ALS-like toxicity. This novel finding enables modeling of neuronal resistance and susceptibility in ALS *in vitro* which could further elucidate mechanisms that could be highly advantageous to activate *in vivo* to rescue vulnerable motor neurons from degeneration.

## EXPERIMENTAL PROCEDURES

### Ethics Statement

All work was carried according to the Code of Ethics of the World Medical Association (Declaration of Helsinki). Animal procedures were approved by the Swedish animal ethics review board (Stockholm Norra Djurförsöksetiska Nämnd). Human CNS samples were obtained from the Netherlands Brain Bank (NBB, www.brainbank.nl), the National Disease Research Interchange (NDRI, www.ndriresource.org) and the NIH Neurobiobank with the written informed consent from the donors or next of kin.

### Generation of mESC lines and specification into oculomotor and spinal motor neurons

Three different mouse ESC lines were used in this study: E14.1 (ATCC), Hb9-GFP (gift from Dr. S. Thams, Karolinska Institutet) and Islet1-GFP (gift from Prof. S. Pfaff, UCSD). To increase oculomotor neuron differentiation efficiency E14.1/NesEPhox2a and Islet1-GFP/NesEPhox2a mESC lines were generated by Lipofectamine (Invitrogen) transfection, NesEPhox2a construct was a gift from Prof. J. Ericson, Karolinska Institutet. As the nestin enhancer was used for overexpression, transgene over-expression is limited to neuronal progenitors. Specification of mESCs into spinal motor neurons was conducted by exposing embryo bodies to the Smoothened Agonist SAG (500 nM, #364590-63-6 Calbiochem) and retinoic acid (RA, 100 nM, #302-79-4 Calbiochem) for 5 days, while oculomotor neurons were patterned by SAG (250 nM) and Fgf8 (100 ng/ml, #423-F8 R&D systems) for 5 days. Media containing morphogens was replaced daily. EB dissociation and motor neuron culture maintenance was carried out as previously described (Wichterle et al., 2002). Oculomotor neurons and spinal motor neurons were kept in Neurobasal media +Glutamax (Gibco) with a cocktail of trophic factor (GDNF, BDNF, NT3, CTNF 5 ng/ml each). All cultures, including survival experiments, were treated with the mitotic inhibitor 5-FDU (final concentration 2μM) to reduce cell proliferation.

### Electrophysiology

Islet1GFP/NesEPhox2A and Hb9GFP cell lines were used to visualize oculomotor and spinal cord motor neurons, respectively. EBs were dissociated and cells were plated onto glass coverslips, coated with Poly-L-ornitine (0.001%, Gibco) and fibronectin (1ug/ml, Gibco), at a density of 200,000 cells per well (24-well plate). The coverslips were placed in a recording chamber and where superfused with with oxygenated recording buffer (95% O_2_/5% CO_2_, 22-24 °C), composed of 111 mM NaCl, 3 mM KCl, 26 mM NaHCO_3_, 1.1 mM KH_2_PO_4_, 2.5 mM CaCl_2_, 1.25 mM MgCl_2_, and 11 mM D-glucose. Patch pipettes (5-6 MΩ) were pulled from borosilicate glass (GC150F-10, Harvard Apparatus) and filled with (in mM): 128 KGlu, 4 NaCl, 10 HEPES, 0.0001 CaCl_2_, 1 glucose, 5 ATP, 0.3 GTP, pH 7.4. Neurons were visualized by video microscopy (Axioskop 2 FS plus, Zeiss) using transversal illumination for increasing contrast and motor neurons were identified under fluorescence using a GFP filter. Whole-cell recordings were performed using a MultiClamp 700A amplifier (Molecular Devices) and acquired using pClamp software (Clampex v.9.2, Molecular Devices). Data were sampled at 20 kHz. Input resistance was measured from the I-V curves; the time constant was fitted with a single exponential curve and the capacity was calculated from the cell time constant and input resistance. The width of the action potential was measured at the half peak of the action potential. Rheobase was defined as the minimal current injection needed to elicit an action potential.

### Fluorescent-Activated Cell Sorting and RNA sequencing of stem-cell derived neurons

Islet1-GFP/NesEPhox2A and Hb9-GFP ESCs were differentiated in oculomotor and spinal motor neurons respectively. GFP+ motor neurons were sorted, and positive and negative fractions were collected for mRNA-sequencing or plated for subsequent immunocytochemistry analysis. FACS was performed using a 100 µm nozzle, a sheath pressure of 15-25 pounds per square inch (psi) and an acquisition rate of 1,000-2,000 events/second (Hedlund et al., 2008). 5×10^3^ cells were collected in a 5% Triton-X-100 (Sigma-Aldrich) in water solution and then prepared for RNA-seq as previously described (Nichterwitz et al., 2016). Samples were sequenced on Illumina HiSeq2000 and HiSeq2500 platforms with readlengths of 43 and 51 bp, respectively.

### Laser capture microdissection and RNA sequencing of human motor neurons

Laser capture microdissection of motor neurons from human post mortem tissues coupled with RNA sequencing, LCM-seq, was performed as previously described (Nichterwitz et al., 2018; Nichterwitz et al., 2016). Briefly, midbrain and spinal cord tissues were sectioned at 10 µm thickness, collected onto MembraneSlide 1.0 PEN microscope slides (Zeiss) and subjected to a quick histological Nissl stain based on the Arcturus Histogene Staining Kit protocol. Slides were then placed under a Leica DM6000R/CTR6500 microscope and captured using the Leica LMD7000 system. Motor neurons were identified by their soma size and their distinct location in axial tissue sections. The cutting outlines were drawn closely around individual cells order to diminish any contamination by surrounding cell. Pools of 20-150 cells were collected and directly lysed in 5µl of 0.2% triton X-100 (Sigma-Aldrich) with RNAse inhibitor (1.5U, Takara). After lysis samples were immediately frozen on dry ice until further processing. Library preparation for sequencing on Illumina HiSeq2000 or HiSeq2500 sequencers was carried out following a modified Smart-seq2 protocol (Nichterwitz et al., 2016).

### RNA-seq data processing

Reads from cell culture samples were mapped to the mouse mm10 reference genome with HISAT2 (Kim et al., 2015), using the publicly available infrastructure (available at usegalaxy.org) on the main Galaxy server (Afgan et al., 2016). Human LCM samples were mapped to hg38/GRCh38 (Gencode version V29) using using STAR version 2.5.3 (Dobin et al., 2013). Transcript-level counts and RPKM values were obtained using rpkmforgenes.py (Ramskold et al., 2009). After mapping, the basic exclusion criterium were <500.000 mapped reads and, for LCM samples, lack of both MN marker transcripts ISL1 and CHAT.

For the human LCM data, independent LCM samples obtained from tissue originating from the same individual (biological replicates) were pooled at the counts level. This ensured that each sample used in analysis was derived from one distinct individual. Ultimately, all 12 spinal cord samples were derived from distinct individuals. Six oculomotor neuron samples originated from distinct individuals, while four more were summed from a total of 12 LCM-seq samples, resulting in a total n=10 samples (from ten individuals) for oculomotor neurons. Two samples were subsequently excluded due to their lack of motor neuron markers, as described above, resulting in a total of eight oculomotor samples. For Onuf’s nucleus, three samples (from three individuals) were used in the analysis, derived from a total of seven LCM-seq samples.

Differential gene expression analysis was performed with the DESeq2 package in R on the raw read counts (version 1.16.1) (Love et al., 2014). Only genes with counts in at least 2 samples were considered for further analysis. No fold change cutoff was implemented and multiple testing correction was performed using the Benjamini & Hochberg correction with an FDR set to 10%. Adjusted p-values < 0.05 were considered significant. Pathway And Gene set OverDispersion Analysis (PAGODA) was run using the SCDE package in R (Fan et al., 2016). All heatmaps were generated in R with either no clustering or *euclidean distance* where appropriate.

### Immunocytochemistry and immunohistochemistry

Coverslips were fixed in 4% paraformaldehyde in 0.1M phosphate buffer (4% PFA) for 20 minutes, washed with PBS and then blocked with blocking solution composed of 10% normal donkey serum or normal goat serum (both from Thermo Fisher Scientific Inc.) and 0.1% Triton-X-100 in PBS for one hour at room temperature. Coverslips were incubated overnight with primary antibodies in blocking solution at 4°C and then incubated with Alexa Fluorophore 488, 568 or 647 secondary antibodies (Invitrogen) one hour at room temperature. The following primary antibodies were used: NF200 (1:1000, #AB5539 Millipore), Tuj1 (1:1000, MRB-435P Covance), Hb9 (1:10, #81.5C10 DSHB), Islet1 (1:500, #ab20670 abcam; 1:100, #40.2D6 DSHB), Phox2A (1:1000, gift of Prof JF Brunet), GFP (1:1000, #ab13970 abcam), phospho-AKT (Ser473) (1:50, #3787 Cell Signaling), activated beta-catenin (1:1000, #05-665 Millipore), alpha-catenin (1:1000, #AB153721 AbCam), ESYT1 (1:100, #HPA016858), BrdU (1:10, #G3G4 DSHB). Hoechst-33342 (Invitrogen) was used as a counterstain. Finally, coverslips were mounted on glass slides using Mowiol 4-88 mounting media (Calbiochem). For immunocytochemistry semi-quantifications and BrdU quantifications at least 3 experiments per condition were counted in technical duplicates. Between 10 and 12 random areas were analyzed per coverslip by blind researcher. For validation of ESYT1 antibody, immunohistochemistry was performed on adult mouse tissue at midbrain and spinal cord levels. Briefly, mice were transcardially perfused with PBS and then with 4% PFA. Brains and spinal cords were dissected and post-fixed for 3 and 1 hour respectively in 4% PFA. Tissue was then cryoprotected with 30% sucrose and then cut at 30 μm thickness. Immunohistochemistry for ESYT1 was performed as indicated above.

### ALS-Like toxicity and motor neuron survival

Oculomotor and spinal motor neurons were cultured for 5 days and then exposed to kainic acid. Three concentrations of kainic acid were tested, 20, 50 and 100 μM. A concentration of 20 μM was found sufficient to induce consistent motor neuron death also when motor neurons were cultured on astrocytes. Kainate toxicity was induced for 7 days. Quantifications were performed blind to the motor neuron population and toxicity condition. Pictures were taken with a Zeiss LSM700 confocal microscope (40x magnification and 1.5 digital zoom) using the ZEN 2010S SP1 software and then further analyzed by ImageJ software. At least four experiments per condition were quantified in technical duplicates. 12 random areas were quantified per each coverslip (at least 130 cells were quantified for each condition). To evaluate proliferation and generation of motor neurons, cultures were exposed to pulses of bromodeoxyuridine (BrdU, #B5002 Sigma) for 24 hours. BrdU was dissolved in Dimethyl Sulfoxide (DMSO, #276855 Sigma) and added to the cultures at a final concentration of 10 μM. After 24 hour exposre to BrdU, cells were washed twice with fresh culture media. Proliferation was assessed at day 0 (D0), day 2 (D2) and day 3 (D3) of the survival protocol and all samples were fixed with 4% PFA at day 7 (D7). Quantifications were performed in triplicates with technical duplicates.

### Morphological analysis

Arborization and Sholl analysis were conducted utilizing Fiji Software. To compare arborization area of OMNs and SC MNs, the microphotographs were transformed to a gray scale, 8-bit image, and quantification was assessed after applying a threshold for signal intensity/background correction. Threshold values were kept constant among images and microphotographs with higher background noise were excluded. Between 10-12 pictures per condition per experiment were quantified. Experiments were performed in quadruplicates. The complexity of oculomotor neuron arborization after KA toxicity was obtained by Sholl analysis (O’Neill et al., 2015) by calculating the average of neurites intersecting defined concentric cycles spaced 25 μm from each other starting from the center of the cell body as a function of distance from soma. Individual oculomotor neurons were analyzed after culturing them at lower density (Spinal motor neuron analysis could not be performed due to their poor survival at lower densities). Several microphotographs were taken at 40x magnification with a Zeiss LSM700 confocal microscope, mounted with Adobe Photoshop software and analyzed with Fiji software. The intersection averages were analyzed by multiple t test.

### Use of published datasets

Microarray data from Kaplan et al. (2014) was obtained through GEO accession number GSE52118.

## ACCESSION NUMBERS

The original RNA sequencing data from this study are available at the NCBI Gene Expression Omnibus (GEO, http://www.ncbi.nlm.nih.gov/geo/) under accession number GSE118620 (*in vitro* generated neuron data) and GSE93939 (human LCM-seq data).

## Statistical Analysis

All statistical analyses were performed with GraphPad software unless otherwise specified. Two-way analysis of variance (ANOVA) was used when taking in account multiple variables and making multiple comparisons, followed by appropriate *post hoc* analysis. Unpaired Student’s t test was used to compare two groups. Further details on statistical tests used can be found in figure legends as well as total number of motor neurons counted per experiment. All experiments were performed at least in triplicates with technical duplicates. All results are expressed as mean ± SEM unless otherwise specified.

## SUPPLEMENTAL INFORMATION

Supplemental information consists of four figures, two tables and can be found within this article online at

## AUTHOR CONTRIBUTIONS

E.H. conceived the study. E.H and I.A. supervised the project. E.H, I.A, J.N., J.A.B., C.S., A.F., and O.K. designed experiments. IA., J.N., J.A.B., C.S., A.F., G.B. and M.C. acquired data. IA., J.N., J.A.B., C.S., A.F., G.B., M.C., O.K and E.H. analyzed data. E.H. wrote the manuscript with the help of I.A. and J.N. All authors edited and gave critical input on the manuscript.

## ACKNOWLEDGEMENTS

This work was supported by grants from the Swedish Research Council (2011-2651, 2016-02112); EU Joint Programme for Neurodegenerative Disease (JPND) (529-2014-7500); Ragnar Söderbergs stiftelse (M245/11), Åhlén-stiftelsen (mA9/h11, mA5/h12, mB8/h13, mB8/h14, mA1/h15, mA1/h16), Hjärnfonden (FO2018-0027), Ulla-Carin Lindquists stiftlese för ALS forskning, and Petrus och Augusta Hedlunds stiftelse (M-2018-0876) to E.H. I.A. was supported by a postdoctoral fellowship from the Swedish Society for Medical Research (SSMF) (2015-2017) and is currently supported by a postdoctoral fellowship from Lundbeck Foundation (2018-2021). J.A.B. is currently supported by a postdoctoral fellowship from SSMF (2017-2019). C.S. was supported by a Early.Postdoc Mobility fellowship from Swiss National Science Foundation (172233). O.K. is supported by Novo Nordisk Foundation Laureate Program.

We would like to thank the Netherlands Brain Bank (NBB, www.brainbank.nl), the National Disease Research Interchange (NDRI, www.ndriresource.org) and the NIH Neurobiobank for providing human CNS samples for this study. We also would like to thank Peter Simonsson for his excellent work in creating the graphical abstract.

