## Supplemental Figures S1-4 for "Modeling motor neuron resilience in ALS using stem cells"

**Supplemental Figure S1. Oculomotor neurons and spinal motor neurons develop electrical properties and mature electrophysiologically in culture.** Cells measured in whole cell configuration in current clamp mode, example traces at day 11 for OMN and SC MN cultures (A; G). Dotted lines indicate 0 mV (A; G). Voltage response to step current injections (-40 pA increasing in 10 pA step) at day 11 in OMNs (A). Increased number of action potentials at day 11 compared to day 9 (B, t test, P= 0.0015, mean±SEM) and reduced width of action potentials (C, t test, P=0.0475, mean±SEM). OMNs at day 11 (n=4) show increased cell capacity (D, t test, P=0.0252, mean±SEM), decreased input resistance (E, t test, P= 0.0270, mean±SEM) and increased rheobase (F, t test, mean±SEM) as compared to Day 9 OMNs (n=9). Voltage response to step current injections (-25 pA increasing in 5 pA steps) at day 11 in SC MN (G). Number of action potentials (H, t test, P=0.0428, mean±SEM) and action potential width (I, t test, P<0.0001, mean±SEM) at day 9 and day 11 in SC MNs. Increased cell capacity (J, t test, P=0.0017, mean±SEM), decreased input resistance (K, t test, P=0.0037, mean±SEM) and rheobase (L, t test, mean±SEM) at day 11 as compared to day 9. ChAT staining in OMNs (M) and SC MNs (N) at day 14 in vitro. ChAT in green, Hb9 in red, Islet1 in blue. Scale bar = 60 μm.

**Supplemental Figure S2. Generation of oculomotor neurons and spinal motor neurons from stem cells and their characterization.** (A) FACS reanalysis plots of Hb9-GFP and (C) Islet1-GFP/NesEPhox2A cells sorted based on GFP expression demonstrated an enrichment of motor neurons (MNs) (Hb9-GFP 85%, Islet-GFP/NesEPhox2A 96% in P3 plots). (B) FAC sorted spinal motor neurons (SC MNs) express Islet1 and Hb9, while oculomotor neurons (OMNs) express Islet1, while lacking Hb9 (D) scale bar = 60 µm. (E) Graph showing mean number of detected genes (and RPKM > 1) (mean ± SEM, t test for RPKM > 0.1 P=0.1196, t test for RPKM > 1 P=0.4047) after mRNA-seq analysis performed on EB dissociation day following FACS of Hb9GFP+ (SC MNs) and Islet1GFP/NesEPhox2A+ (OMNs) cells. Both conditions show similar amounts of expressed genes. (F) Heatmap showing the differential Hox gene expression between OMNs and SC MNs, confirming their positional identity. (G) DESEQ analysis performed on transcripts obtained from generated oculomotor neurons (OMNs) and spinal motor neurons (SC MNs) reveal significant differences in genes controlling axon guidance, $Log_{10}$-transformed data for *Sema6d*, *Plxa4*, *Cdh6* and *Cdh12*, preferentially expressed in OMNs, and *Sema4a*, *Sema5b*, *Epha3* and *Ephx4* preferentially expressed in SC MNs (adjusted *P < 0.05). (H) Preferntial expression of Sema6d in oculomotor neurons in the published Kaplan et al data set (t test, *P* <0,001). (I) Graph indicated percentage of BrdU+ cells over Islet1+ cells at D7 assay. BrdU pulses were performed at D0, D2 and D3 survival assay quantifications were performed at D7. (mean ± SEM, 2way ANOVA and Tukey's multiple comparison test, F(2, 169)=0.3278, P=0.7209, n=3). 10 random areas were quantified per each time point per experiment (in duplicates), with at least 100 cells quantified per condition.

**Supplemental Figure S3. Preferential expression of Ca-regulating genes in oculomotor neurons.** (A) After DESEQ analysis of generated oculomotor neuron (OMN) and spinal motor neuron (SC MNs) mRNA expression, several transcripts of $Ca^{2+}$ binding proteins could be found differentially expressed. Graph shows four of them (*Cald1*, *Esyt1*, *Camk2a*, *Hpcal1*) with preferential expression in OMNs suggesting increased capability of intracellular $Ca^{2+}$ buffering during excitotoxicity (adjusted *P<0.05, $log_{10}$-transformed RPKM values). (B-C) ESYT1 immunohistochemistry performed on OMNs and SC MNs at D7 toxicity assay revealed preferential expression in OMNs at protein level. Scale bar = 60 µm. (D) Semi-quantification of ESYT1 staining in control and KA20 conditions (mean ± SEM, 2way ANOVA and Tukey's multiple comparison test, F(1, 239)=31,84, *** P < 0.0001, n=3). Experiments were performed with technical replicates and with at least 100 motor neurons counted per condition.

**Supplemental Figure S4. LCM-seq of human resilient and vulnerable motor neuron groups.**

Laser capture microdissection (LCM) was used to isolated oculomotor neurons (A-D), spinal (cervical and lumbar) motor neurons (E-H) and Onuf's nucleus motor neurons (I-L) from human *post mortem* tissues.

RNA sequencing of isolated neurons showed that (M) motor neurons also expressed Islet-1/2 and ChAT, while being almost devoid of contaminating glial markers. (N) Analysis of the Phox2a/b and Hox gene expression clustered oculomotor neurons away from all spinal motor neuron groups. (O) Analysis of the PI3K-AKT signaling pathway.

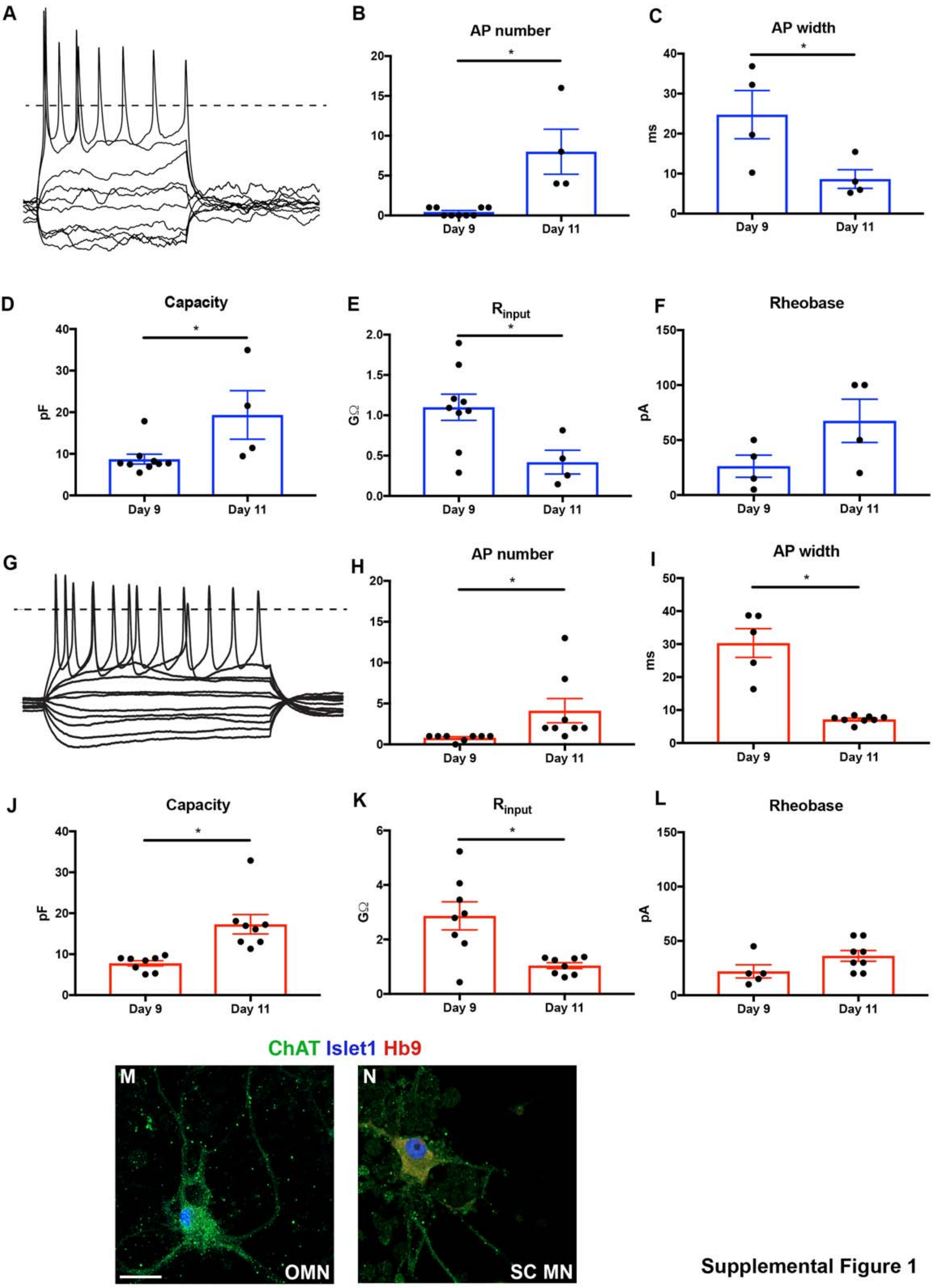


Supplemental Figure 1

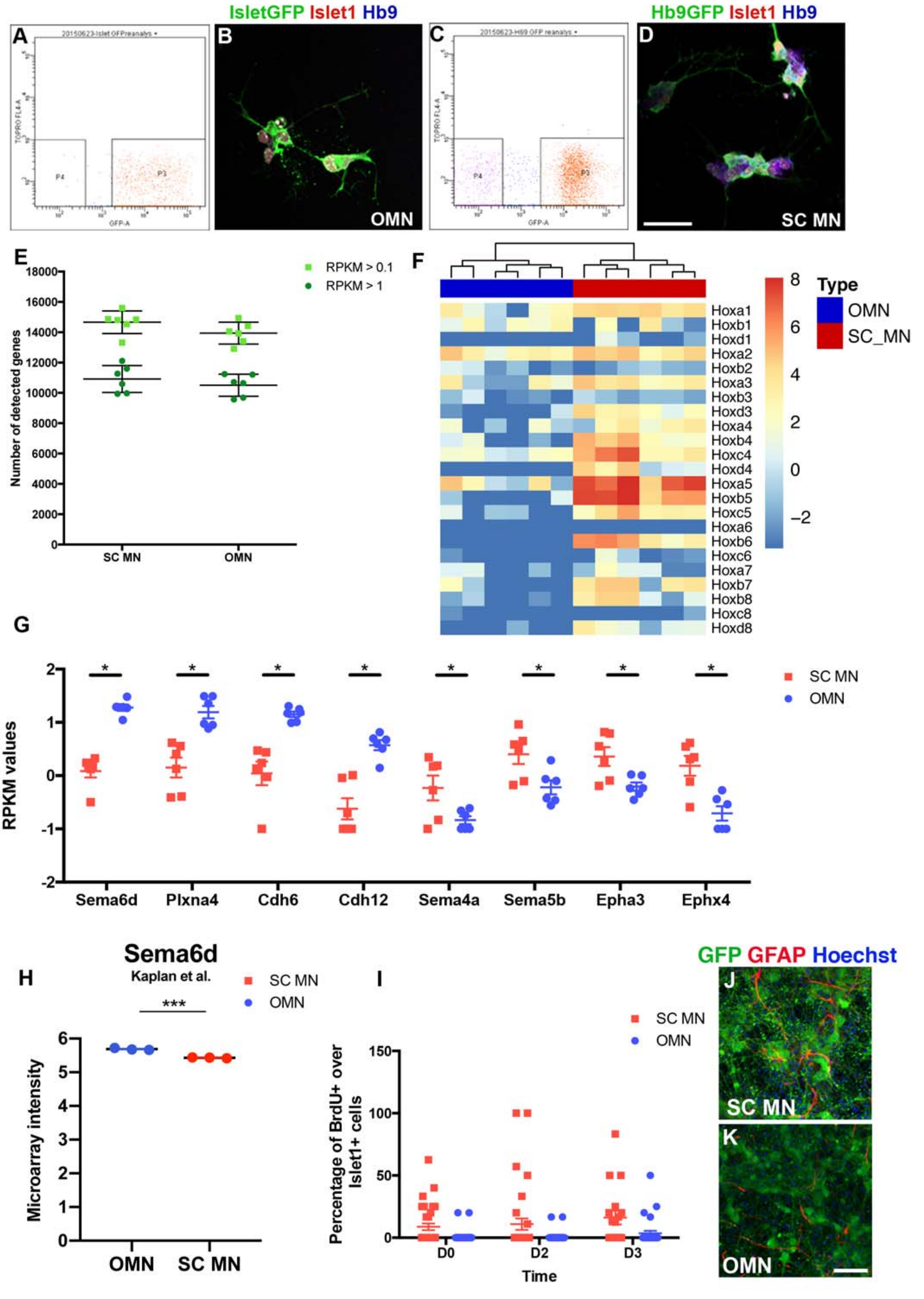


Supplemental Figure 2

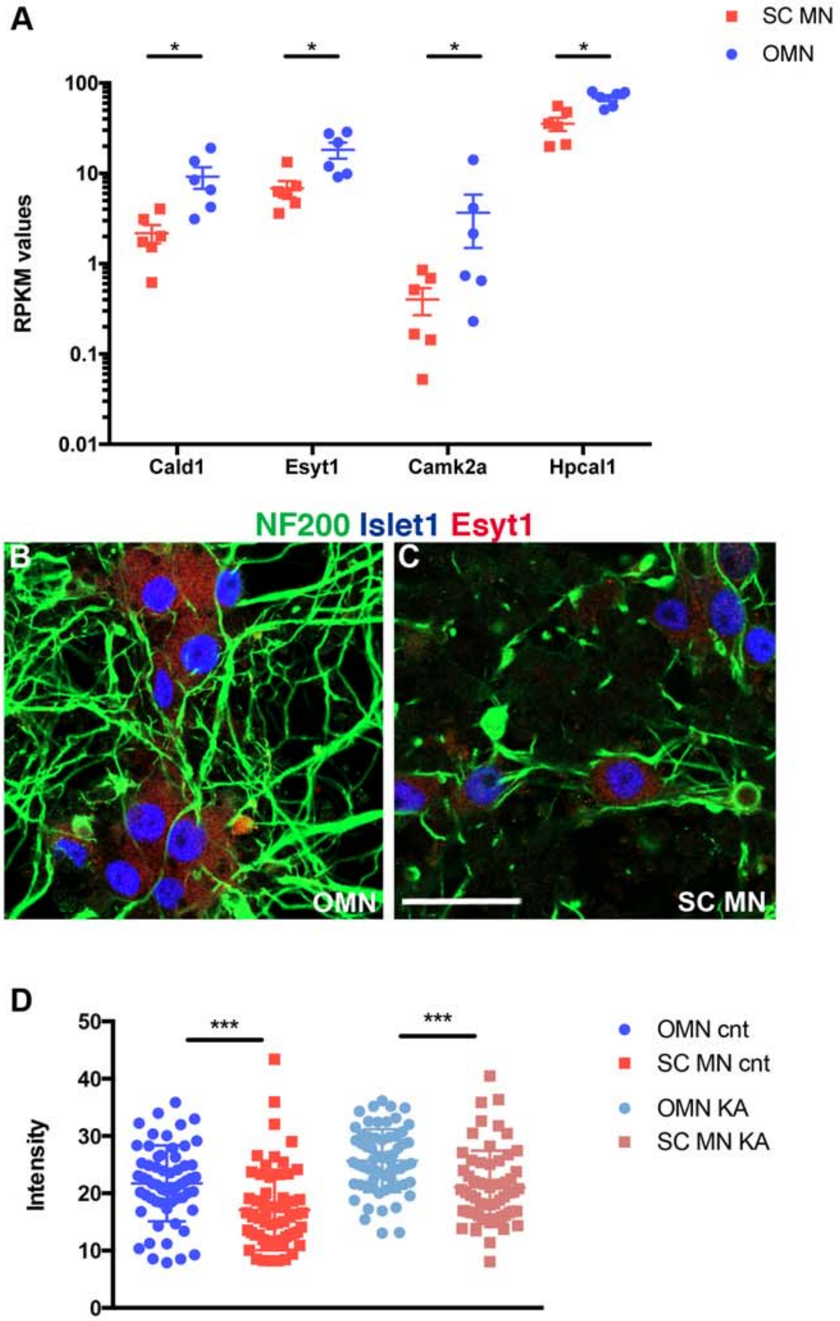


Supplemental Figure 3

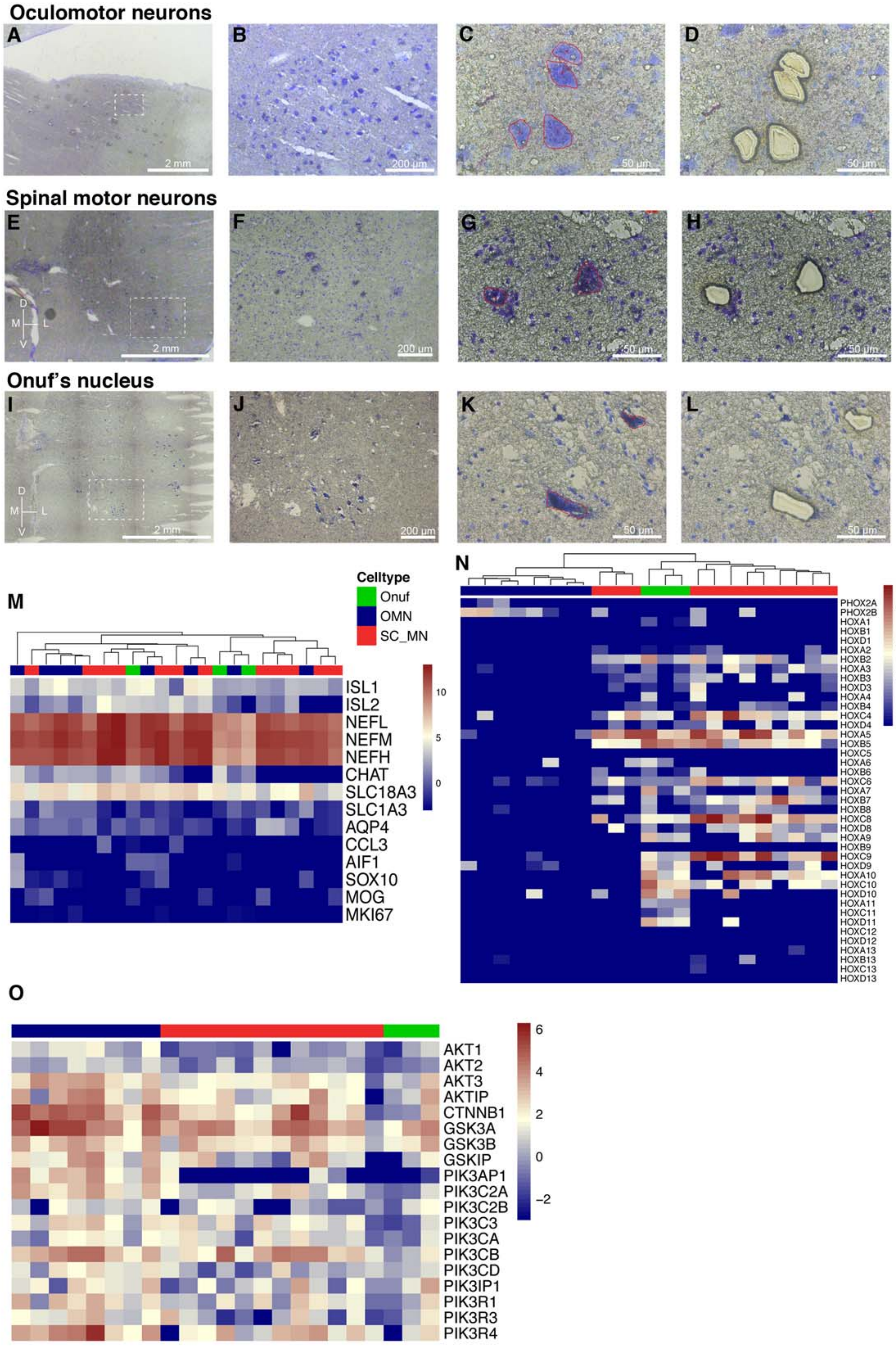


**Supplemental Figure S4**
